# A Human Accelerated Region participates in early human forebrain patterning and expansion

**DOI:** 10.1101/777235

**Authors:** Sandra Acosta, Jaydeep Sidhaye, Luciano Fiore, Isabel Rollan, Giovanni Iacono, Alexander V. Misharin, Nozomu Takata, Martin Sikora, Nikita Joshi, Beisi Xu, Eske Willerslev, Holger Heyn, Miguel Manzanares, Juergen A. Knoblich, Guillermo Oliver

**Affiliations:** Center for Vascular and Developmental Biology, Feinberg Cardiovascular Research Institute, Northwestern University, Chicago, IL; Institute of Molecular Biotechnology of the Austrian academy of sciences (IMBA), Vienna, Austria; Centro Nacional de Investigaciones Cardiovasculares (CNIC), Madrid, Spain; CNAG-CRG, Centre for Genomic regulation (CRG), Barcelona Institute of Science and Technology (BIST), Barcelona, Spain; Division of Pulmonary and Critical Care Medicine, Northwestern University, Chicago, IL; GeoGenetics Center, University of Copenhagen, Copenhagen, Denmark; Bioinformatic Department, St. Jude Research Children’s Hospital, Memphis, Tn; Universitat Pompeu Fabra (UPF), Barcelona, Spain; Centro de Biologia Molecular Severo Ochoa, CSIC-UAM, Madrid, Spain

**Keywords:** Human brain evolution, Forebrain organoids, HAR, Enhancer regulation, Cerebral Cortex, scRNAseq, ESC

## Abstract

The expansion of the mammalian brain is associated with specific developmental processes; however, not much is known about how evolutionary changes participated in the acquisition of human brain traits during early developmental stages. Here we investigated whether enhancers active during the phylotypic stage show human-specific genomic divergence which could contribute to the evolutionary expansion of the forebrain. Notably, we identified an active enhancer containing a human accelerated region (HAR) located in the Chromosome 14q12, a region enriched with neurodevelopmental genes, such as Foxg1, Nkx2.1 and Nova1. Reporter analysis revealed that the human variant is active in the forebrain in transgenic mice and that it has stronger enhancer activity than the mouse or chimpanzee versions. Humanization of the mouse enhancer variant in transgenic mice and in mouse organoids resulted in an expansion of Foxg1 expressing domains in the forebrain early neural progenitors with a bias towards dorsal identities. Overall, our results suggest that human-specific mutations in critical regulatory elements controlling early brain development impact the expansion and patterning of the forebrain.

## Introduction

The evolution of the forebrain is responsible for the acquisition of human higher cognitive functions. Most of these gained extraordinary evolutionary abilities lay on the cerebral cortex, the largest and the anterior-most structure of the brain (Hill & Walsh, 2005). The human cerebral cortex and its developmental precursor, the forebrain, have undergone rapid evolutionary expansion in size and cellular diversity (Hill & Walsh, 2005; Lui *et al*, 2011). Therefore, investigating the human-specific molecular mechanisms that contribute to the evolution of the human forebrain is a central question towards understanding human brain development and the acquisition of human-most cognitive traits.

Forebrain development is a stepwise process spanning from the initial patterning of the neuroectoderm during late gastrulation, to the formation of cerebral cortex circuits postnatally. Despite the obvious morphological differences that the vertebrate forebrain exhibits during late embryogenesis, they are almost indistinguishable during the earliest developmental stages. These remarkable initial resemblances during the phylotypic stage among very diverse organisms, are partially a consequence of the high level of conservation of the transcriptional networks controlling embryonic patterning and differentiation (Davidson & Erwin, 2006; Wilson & Houart, 2004; De Robertis, 2008; Duboule, 1994; Irie, 2017). In the case of the central nervous system (CNS), despite the limited divergence in genes controlling neural development, their spatio-temporal pattern of expression varies amongst species, suggesting that evolution-driven changes in the activity of enhancers play a central role in brain divergence in the animal kingdom (Sylvester *et al*, 2010). Indeed, enhancers are more susceptible to mutations than genes, and they seem to have a prominent effect in the evolution of the human brain (Reilly & Noonan, 2016; Bae *et al*, 2014; Franchini & Pollard, 2015; 2017).

Extensive whole genome comparative studies performed in the last few years lead to the identification of Human Accelerated Regions (HAR) as putative drivers of human-specific traits (Pollard *et al*, 2006; Boyd *et al*, 2015; Doan *et al*, 2016; Prabhakar *et al*, 2006). HAR are highly conserved mammalian genomic regions that acquired mutations in humans. Moreover, the presence of HAR substitutions in enhancers involved in human neural development have been associated with the gain of human traits such as the presence of an expanded cerebral cortex (Boyd *et al*, 2015), or the impairment in cognitive functions in a variety of neurological disorders when mutated (Doan *et al*, 2016). However, little is known about the presence and roles of HAR in regulatory elements active during early neural development and their potential impact on cerebral cortex formation.

In this study, we explored the presence of HAR in mouse enhancers active during the gastrula to neurula transition stages. We report on the identification of a HAR enhancer located in a genomic region enriched in neurodevelopmental genes. Analysis on genetically modified transgenic mice and embryonic stem cells (ESCs)-derived organoids harboring the human enhancer variant suggest a central role of this regulatory element in brain development, and in enhancing forebrain neuroepithelium proliferation and patterning. In humans, further layers of genomic regulation seem to play a role in the activity of this enhancer, since chemical knock-down or heterozygous mutations do not have a phenotypically identifiable effect. This study shows for the first time that some evolutionary changes that could lead to human-specific brain traits are acquired during early embryonic stages.

## Results

### Identification of human-specific evolutionarily changes in enhancers active during early forebrain development

During the phylotypic stage, around neurulation, most vertebrates look very similar (Duboule, 1994). However, not much data is yet available about critical evolutionary changes impacting early neurodevelopmental genes during the transition from gastrula to neurula. This lack of knowledge is likely due to the fact that most processes involved in early development are highly conserved during evolution (Dickel *et al*, 2018); therefore, even subtle changes in their early expression could have a big impact later in life. Thus, we decided to search for putative evolutionary mutations in enhancers active during early embryonic stages (late gastrula-early neurula) we performed H3K27ac ChIP-seq in E7.5-8.0 mouse embryos (late gastrula to neurula) (Creyghton *et al*, 2010; Heinz *et al*, 2015). Four replicates containing 25-30 pooled embryos each were prepared and bioinformatically analyzed (Fig 1A). Only peaks consistently present in the replicates were considered for further analysis. To determine whether any of the identified active enhancers could be involved in the acquisition of human evolutionary traits at this early stage of neural patterning, the H3K27ac ChIP sequencing database was cross-compared with previously published databases for human accelerated regions (Fig 1A) (Pollard *et al*, 2006; Prabhakar *et al*, 2006). This analysis identified a total of 655 active HAR-containing genomic regions during late gastrula-neurula. Mapping annotation of these HAR indicated that 101 were located nearby (up to 1 Mb) developmental genes. Out of those, 10 were in the vicinity of known neural development genes (Fig 1B). Amongst those, of particular interest was a 795 bp HAR located in Chromosome 14q12, a region coding for neurodevelopmental genes such as Foxg1, NOVA1, PRKD1 and NKX2.1. Notably this region is located 630 kb upstream of Foxg1 (Fig 1C), a gene critical for forebrain development, whose expression starts around E8-8.5 in the mouse embryo (Xuan *et al*, 1995; Geng *et al*, 2016). This HAR enhancer, herein HAR-14q12, is highly conserved amongst mammals, i.e. only 8 nucleotides are accelerated mutations in humans (Fig 1C, red colored lines), sharing more than 90% homology with chimps and gorillas. When compared to the mouse variant (mHAR-14q12), it only diverged in 18 nucleotides (17 single nucleotides and one insertion) (Fig S1).

**Fig 1.**
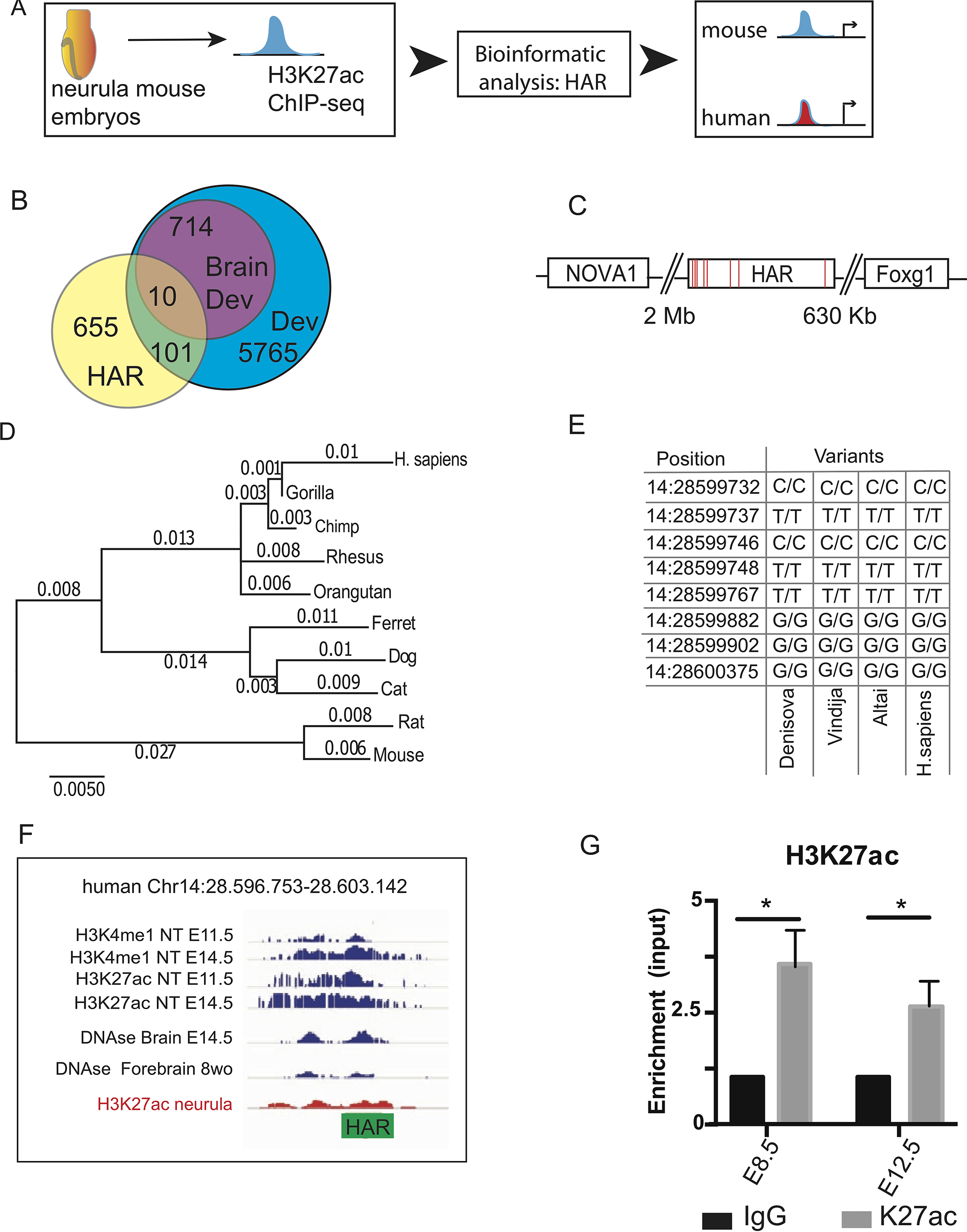
HAR enhancer is active during early development. A. Workflow of the study showing the initial H3K27ac ChIP-seq performed on late gastrula-early neurula stage (E7.5-E8) mouse embryos to identify active enhancers. Bioinformatics analysis was performed to explore the presence of HAR elements in the enhancers after LiftOver processing to convert the mouse genome into human. B. Venn diagram showing the number of HAR identified in active developmental enhancers, from which 10 where located in the vicinity of neural development genes. C. Scheme of the identified hHAR region located flanked by *Foxg1* and *Nova1*. Red lines indicate the positions of the Human-accelerated substitutions. D. Maximum-likelyhood phylogenic tree for the orthologous mammalian loci’s. E. Matrix indicating the nucleotides accelerated in *H. sapiens* compared to Neanderthals (Altai and Vindija) and Denisovans (Denisova) genomes. F. Bioinformatic analysis of the presence of acetylated peaks upstream of *Foxg1*. This enhancer shows regulatory activity during different developmental stages in various neural tissues (NT), including the generated late gastrula-early neurula database (in red). G. H3K27ac ChIP analysis showing that HAR-Foxg1 exhibits enhancer activity during forebrain development (E8.5 foreheads and E12.5 forebrains).

Maximum likelihood estimation on the 795 bp full-length enhancer indicates a clear divergence in primates with respect to other clades, with dramatic acceleration only in the human lineage (Fig 1D). Notably, once the human lineage diverged from the other apes, 8 accelerated single nucleotide substitutions were rapidly acquired, as indicated by the increased length of the human branch in the maximum likelihood estimation (Fig 1D). Moreover, comparative analysis between *H.neanderthal* (Vindija and Altai), *H.denisovan* (Denisova) and modern humans showed 100% identity in the enhancer sequence, suggesting that the accelerated nucleotide substitutions found in HAR-14q12 were fixed before the divergence from a common ancestor (Fig 1E). The accelerated nucleotide substitutions in the HAR region resulted in substantial changes in the combination of transcription factor binding sites (TFBS), possibly providing the enhancer with putative new functions (Fig S1).

To further delve into the functional role of the enhancer, we conducted a meta-analysis of the epigenomic signature of this region to derive information about the developmental stage and the tissues where this enhancer is likely active. Such analysis was performed taking advantage of publicly available epigenomic databases (see Materials and Methods) for regulatory elements (H3K4me, H3K27ac, DNAse I hypersensitivity) at different stages of neural development in the neural tube, the forebrain and the retina in the mouse and in human cells. The combined analysis suggested that HAR-14q12 is active throughout early neural differentiation stages (Fig 1F). Accordingly, we further analyzed the activity of this enhancer in the forebrain at relevant developmental stages: E8.5 (neural tube closure and onset of Foxg1 expression) and E12.5 (early neurogenesis and rapid expansion of telencephalic progenitors). The analysis of ten E8.5 foreheads and five E12.5 forebrains (pooled together in triplicates) by H3K27ac ChIP and qPCR confirmed the *in silico* evidence about HAR-14q12 enhancer activity during critical steps of forebrain development (Fig 1G). Indeed, these results suggest that HAR-14q12 is an active enhancer that spans its activity throughout early forebrain development, including the neurula stage, once Foxg1 starts to be expressed.

### HAR-14q12 is an active enhancer in the central nervous system

Next, we addressed whether HAR-14q12 was an active enhancer *in vivo* in the developing CNS. To do this, we inserted the full length (795bp) HAR-14q12 into a LacZ reporter vector, upstream of the hsp68 minimal promoter (Pennacchio *et al*, 2006). Transient transgenic embryos were generated by pronuclear injections. Following X-gal staining, ß-galactosidase activity was detected in the forebrain of 13/23 E9.5 embryos (Fig 2A-C). As seen in Fig 2, X-gal staining or ß-galactosidase immunostaining revealed that HAR-14q12 activity was mostly restricted to neural tissue, particularly along the ventricular zone of the forebrain (Fig 2A-I and Fig EV1A-B). To determine whether at E9.5 the HAR-14q12 enhancer activity overlapped with endogenous mouse Foxg1 expression, we performed ß-galactosidase immunostaining on top of Foxg1 RNA *in situ* hybridization in sagittal sections. As seen in Fig 2D-F, endogenous Foxg1 was localized as expected in the telencephalic region for the forebrain; however, HAR-Foxg1 expression appears expanded along the entire forebrain, including a region of high ß-galactosidase staining in the ventral diencephalon where endogenous mouse Foxg1 is not expressed at this stage (Fig 2I and Fig EV1A-B). The H3K27ac ChIP indicated that the enhancer was still active in the mouse forebrain at E12.5 (Fig 1G). Thus, we explored whether the human enhancer variant was also active at E12.5 in the forebrain in HAR-LacZ transgenic embryos. X-gal staining was detected in the dorsal and ventral forebrain, as well as in the eye and the ventral diencephalon at this stage.

**Fig 2.**
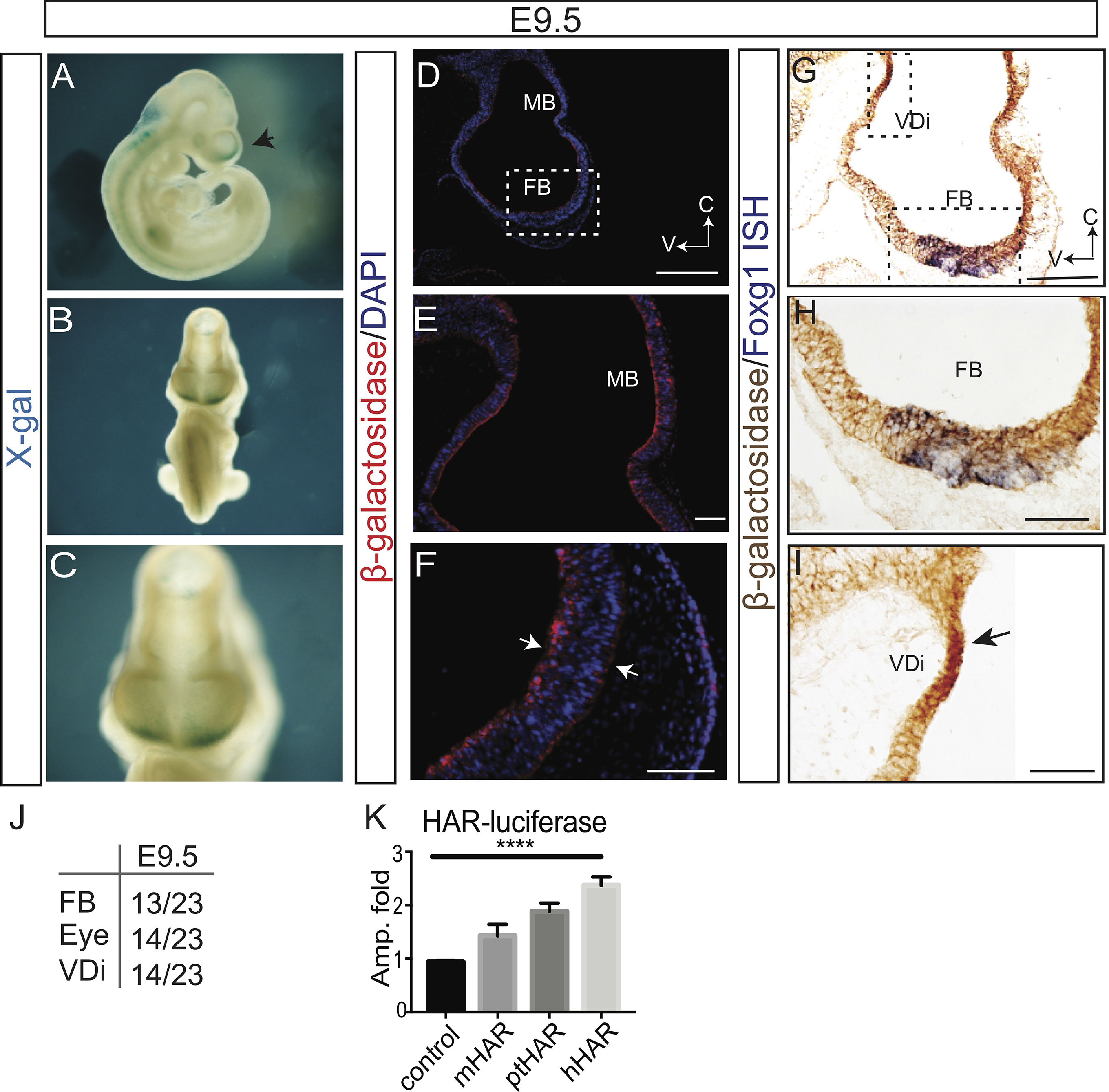
HAR-Foxg1 exhibits expanded enhancer activity in the developing forebrain. A-C. hHAR-LacZ transgenic E9.5 embryos stained for X-gal showing expression in the telencephalic vesicles. The arrow shows the stained telencephalic vesicle. Frontal view of the embryo (C) showing staining in the most anterior region of the telencephalic vesicles, namely the ANR. D-F. β- galactosidase immunostaining in sections of E9.5 hHAR-LacZ transient transgenic embryos. β-galactosidase is detected in the neuroepithelium of the entire forebrain. The inset (F) is a higher magnification of the anterior forebrain (FB) region shown in D that matches the endogenous expression at this stage shown in red in D. Arrows indicates the apical and basal processes of the radial glia stained with β-galactosidase. G-I. β- galactosidase immunostaining together with Foxg1 *in situ hybridization* on E9.5 transgenic embryos. Boxes in G correspond to the higher magnifications shown in H and I. G Overlap between endogenous Foxg1 expression and b-galactosidase is seen in the forebrain. I. Strong levels of b-galactosidase (black arrow) expression are seen in the ventral diencephalon (VDi) where endogenous Foxg1 expression is not present. J. Number of embryos showing X-gal staining in the forebrain, eye and ventral diencephalon. K. Luciferase analysis indicating a significant increase in activity with the HAR-14q12 enhancer in comparison with the mouse (m), the chimp (pt) and the human (h) variants. This experiment was performed twice and in quadruplicates (ANOVA, error bars show s.e.m). Scale bars A,D: 200mm and B,C,E,F:100mm

This initial analysis argued that at E9.5, the HAR-14q12 variant drives activity into the entire mouse developing forebrain. Furthermore, bioinformatics analysis showed that in humans these accelerated substitutions promoted a few changes in the TFBS predicted to bind to the hHAR-14q12 region. This is evident when comparing the human and chimp variants, and even more when compared to the mouse variant (Fig S1). Accordingly, these changes in TFBS promoted in the human enhancer variant could lead to differential regulatory activity amongst species.

It has already been shown that slight alterations in enhancer activity can have a dramatic phenotypic impact in the differentiation and function of the brain (Sylvester *et al*, 2010). In addition to promoting spatio-temporal changes in the regulatory activity of enhancers, mutations in TFBS might also impact their regulatory strength (Grossman *et al*, 2017). To evaluate whether this could also be valid for HAR-14q12, we inserted the full-length mouse (mHAR), chimp (ptHAR) and human (hHAR) enhancer variants into a luciferase vector. As seen in Fig 2K, the highest luciferase activity was detected under the regulation of the hHAR, followed by the chimp variant and the mHAR was the one with the lowest activity. The observed differences in luciferase activity might be related to the number of nucleotide substitutions present in each of those HAR. Indeed, while mHAR and hHAR diverged in 18 out 795 nucleotides, the chimp only diverged in 8 nucleotides with the hHAR. Interestingly, 7 out of the 8 HAR nucleotide substitutions are located very close to each other in the 5’ region of the enhancer (Fig EV1G). Luciferase analysis using the 5’ short human HAR-14q12 fragment significantly lowered the activity compared to the full-length fragment (Fig EV1H), suggesting that the whole fragment is necessary for proper gene regulation. In summary, these results suggested that in the embryonic forebrain, the accelerated substitutions in HAR-14q12 might not only affect the spatial expression of the regulated gene, but also impact its expression levels in those tissues.

### Human enhancer variant promotes forebrain neuroepithelium expansion in transgenic mice

In an effort to further determine whether the evolutionary acceleration of the human enhancer variant had an impact in forebrain development and in the regulation of Foxg1, we tested the putative differential activity of the mouse and human variants. Using a vector with Foxg1 fused to GFP (Baek *et al*, 2015) under the regulation of either the mouse or the human enhancer variants transgenic mice were generated (Fig 3A). At E9.5, while the mouse enhancer variant produces normal embryos, those injected with the human variant showed high mortality and significant forehead malformations (Fig EV2A-B). GFP immunostaining did not show significant differences between the expression domains of the mouse and the human enhancer variants, suggesting that they are both active in similar brain regions. However, the area expressing endogenous Foxg1 and Nestin were expanded in the human variant *vs* the mouse variant (Fig 3B-D, H and Fig EV2C-D), suggesting that changes in the patterning of the forebrain begun at early neurula stages when the neuroepithelium started expanding. Next, pH3 immunostaining suggested that this expansion was due to an increase in the proliferation of neuroepithelium progenitors (Fig 3E-G and I), rather than a change in their identity. Altogether, the human enhancer variant seems to play a role in forebrain patterning by increasing the pool of progenitors starting at neurulation.

**Fig 3.**
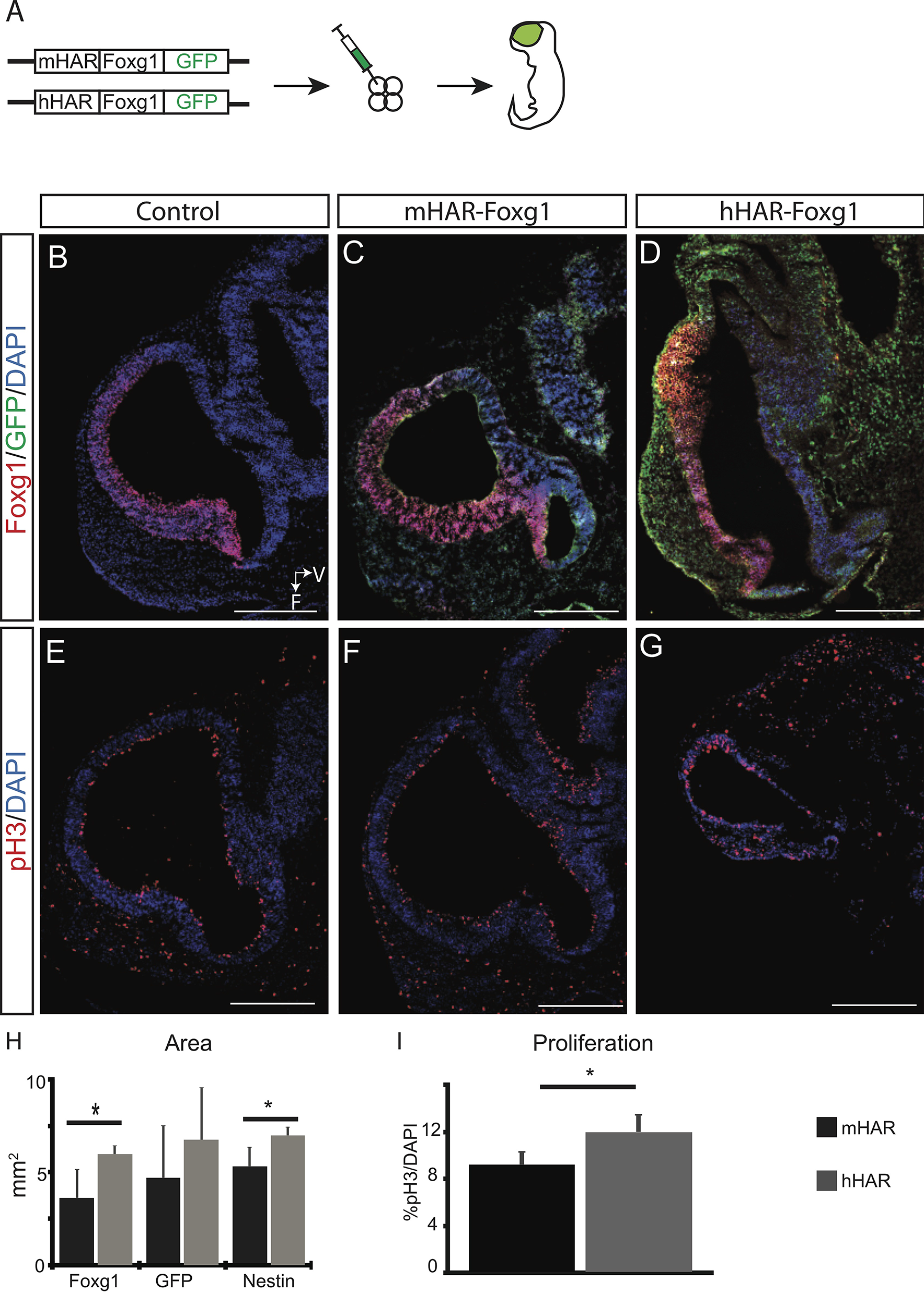
Human enhancer variant drives stronger Foxg1 expression than the mouse enhancer variant in the forebrain of transgenic mice. A. Scheme of the constructs used to generate transgenic embryos carrying the Foxg1-GFP fusion protein regulated by the mouse or the human enhancer variants. B-D. E9.5 transgenic forebrain stained against endogenous Foxg1, GFP and DAPI in WT controls. mHAR-Foxg1 and hHAR-Foxg1 show an expanded but disrupted forebrain territory. E-G. pH3 staining showing dividing progenitors. H. Measurement of the area stained with endogenous Foxg1, GFP as a marker of transgenic Foxg1 and Nestin. I. Percentage of dividing pH3+ progenitors over the total cells (DAPI) in the neuroepithelium. Scale bars: 500mm

### HAR-14q12 increases Foxg1 expression in humanized forebrain organoids

The results above indicate that the evolutionary changes in the human enhancer variant potentiate its regulatory activity during early forebrain development. Next, we wondered if it regulates Foxg1 expression and thus, whether is involved with the expansion of the human forebrain neuroepithelium. Indeed, overexpression of Foxg1 has been associated with an increase in the number of neuroblasts during chick embryogenesis (Ahlgren *et al*, 2003). To this purpose, we took advantage of an available Foxg1-Venus reporter mouse ESC line (Eiraku *et al*, 2008) to generate ESC-derived 3D forebrain organoids that recapitulate key features of early forebrain development (Nasu *et al*, 2012). To do this, we first generated a heterozygous mHAR knock-out by introducing indels using CRISPR/Cas9 (Fig EV3B). This approach allowed us to evaluate the effect of the mouse variant in cis, through changes in the expression of the Foxg1-Venus reporter. Additionally, we targeted the hHAR variant into the other allele to evaluate possible negative effects in mouse forebrain differentiation and growth promoted by the human variant. Using this mHAR KO/hHAR reporter mouse ESC line we generated mouse forebrain organoids (Fig EV3A). Compared to control organoids, most targeted organoids (n=12 per two clones per two experiments) grown from independently generated clones showed a significant decrease in Venus expression, while no significant changes were observed in the acquisition of forebrain identity as indicated by Six3 and Pax6 expression (Fig EV3C-J). These results confirmed that the mHAR locus regulates expression of Foxg1.

Nevertheless, if the enhancer is involved in forebrain expansion in humans, this phenotype should be observed using a targeted gain-of-function approach. Accordingly, we knocked-in the human HAR variant into the orthologous HAR locus to generate humanized mouse ESCs (hHAR-ESC) using a novel CRISPR/Cas9 strategy (Acosta *et al*, 2018) (Fig 4A). Using this approach, off-targets were not detected in the hHAR-ESCs upon CRISPR/Cas9 treatment. Following the 3D forebrain organoids differentiation protocol (Fig EV3A), Foxg1 expression was stronger and expanded in the day 7 “humanized” hHAR-ESC-derived organoids when compared with their isogenic controls (day 7 is the earliest stage when Foxg1 expression is detected in controls) (Fig 4B,C, F). Although by day 10 most cells in control organoids expressed Foxg1 (Fig 4D), the levels of Foxg1 expression remained higher (Fig 4E-G) in the humanized hHAR-ESC-derived organoids.

**Fig 4.**
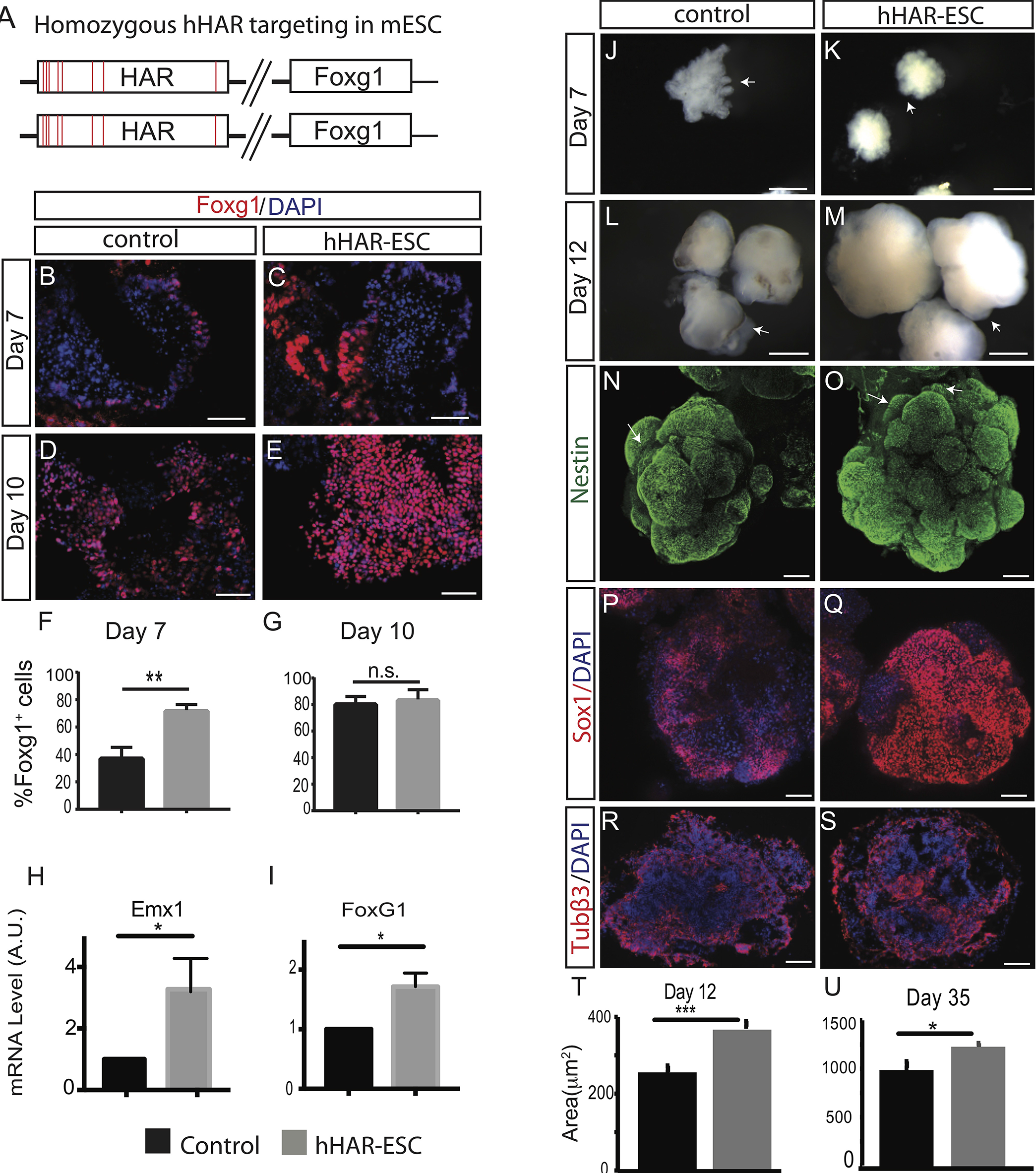
HAR-14q12 induce progenitors expansion in mouse ESCs. A. Homozygous knock-in strategy of the human HAR-Foxg1 enhancer variant in mouse ESCs. B-G. Increase in endogenous Foxg1 expression in humanized ESCs containing the HAR enhancer (hHAR-ESC). An increase in Foxg1 levels is already seen at day 7 (early neurogenesis) in the humanized organoids (B,C). By day 10 (D, E) the number of Foxg1 expressing progenitors has largely increased in the humanized organoids. (F, G) Quantification of Foxg1 stained cells in neural vesicles at day 7 (F) and day 10 (G) of differentiation (n=at least 3 aggregates per triplicate from 2 independently obtained targeted clones). ** p<0.01. H,I. mRNA gene expression analysis in day 7 organoids indicating an increase of Foxg1 (H) and Emx1 (I) expression. J-S. Humanized hHAR-ESCs organoids (K,M) are larger than controls (J,L) as the differentiation proceeds. This increase is followed by an increase in the number of Nestin^+^ (N,O) and Sox1^+^ (P,Q) progenitors in the humanized mESCs. R,S. The amount of differentiated neurons is not obviously increased in hHAR-ESCs at day 12. Measurement of the area showing a significant increase at day 12 (T) and day 35 (U) of differentiation. Statistical significance was calculated with T-Test. Scale bars B-F: 50mm L-S:150mm.

These results suggest that the evolutionary changes induced by the human enhancer variant are responsible for an increase in the number of Foxg1 expressing cells at early neural stages when the neuroepithelium is still expanding.

### HAR-14q12 is responsible for the expansion of neural progenitors

We next addressed the consequences of expanding Foxg1 expression during early developmental stages using the “humanized” hHAR-ESC organoids. In animal models, Foxg1 is a transcriptional repressor that plays an important role in the expansion of forebrain neural progenitors (Xuan *et al*, 1995; Yao *et al*, 2001) and when expressed ectopically, it expands the neural progenitor pool (Ahlgren *et al*, 2003) supporting its essential role in neural induction. To assess the effect of differential Foxg1 regulation by the mouse and the human HAR variants, three independent clones of the isogenic hHAR-ESCs and controls were seeded and differentiated into forebrain organoids up to 35 days (Nasu *et al*, 2012). The targeted humanization of the enhancer variant did not produce changes in cell cycle in the undifferentiated hHAR-ESCs (Fig. S2). At day 7, Foxg1 and Emx1 expression levels were increased in the humanized; however, no obvious morphological changes in the size and shape of the organoids were observed at this stage (Fig 4H-K). Changes became evident at day 12 of differentiation, a stage when radial glia is expanding and neurogenesis has started. At this stage, we observed that neural progenitors increased dramatically leading to a significant increase in the hHAR-ESC organoid size (Fig 4L-U).

To further investigate the mechanisms responsible for the increase in the organoid size, we focused on day 12 organoids, the stage when size differences were more evident (Fig 4L,M) and when the neuroepithelium is already committed to forebrain and radial glia (RG) fates and continues expanding. Whole-mount immunostaining of day 12 organoids showed that most, if not all cells in the larger size hHAR-ESC derived organoids were neural progenitors, as indicated by the expression of Nestin and Sox1 (Fig 4N-Q). Notably, the increase in the number of progenitors was accompanied by an increase in the amount of vesicle-like structures in the organoids’ surface (Fig 4N, O). Moreover, as indicated by ß3-tubulin immunostaining, the increase in neural progenitors did not result in an obvious expansion in the number of neurons at this stage (Fig 4R, S). Even at day 35, when neurogenesis is well established, humanized hHAR-ESC organoids contained an increased number of neural progenitors than controls (Fig S3A-D). Moreover, at this late stage the levels of neurogenesis and gliogenesis were also higher in hHAR-ESC organoids than in the controls (Fig S3C-F). Altogether, these results suggest that the presence of the human enhancer variant in the organoids allows progenitors to remain longer in a proliferative stage without altering their maturation into neurons and glia.

### Progenitors derived from hHAR-ESCs cycle faster than controls

Next, we investigated the mechanisms that potentially lead to an increase in the number of progenitors and increased tissue size in hHAR ESC-derived organoids by day 10-12. Foxg1 expression has been associated with the control of cell proliferation and apoptosis (Manuel *et al*, 2010; Seoane *et al*, 2004). Hence, we assessed for cell cycle and apoptosis differences in the organoids at day 7, soon after the start of Foxg1 expression (Fig 4B-D) and at day 10, when neural progenitors expansion is at its highest rate. (Fig 5). Upon immunostaining with the mitotic marker phospho-Histone 3 (pH3), we observed that at both developmental stages the number of mitotic cells was significantly higher in the hHAR-ESC-derived organoids (Fig 5A-E). Next, we asked if this increase in mitotic cells was due to faster cell cycle. To test this, we measured the mitotic index, a cell cycle length measurement, by performing a pulse of EdU at day 10, followed by pH3 staining of organoids harvested after 2 and 6 hours. The mitotic index, measured as the number of cells double labelled for pH3 and EdU informs about the cells that divided recently. The faster the cell cycle, the larger the number of cells that will be double labeled at the second time point. Indeed, we observed that the number of pH3 and EdU double labelled cells located in the vesicle-like structures that contain Nestin^+^ neural progenitors was significantly higher in the hHAR-ESC derived organoids after 6 hour incubation (Fig 5 C,D,F). These data confirmed that the neural progenitors in the hHAR-ESC organoids proliferate faster than controls.

**Fig 5.**
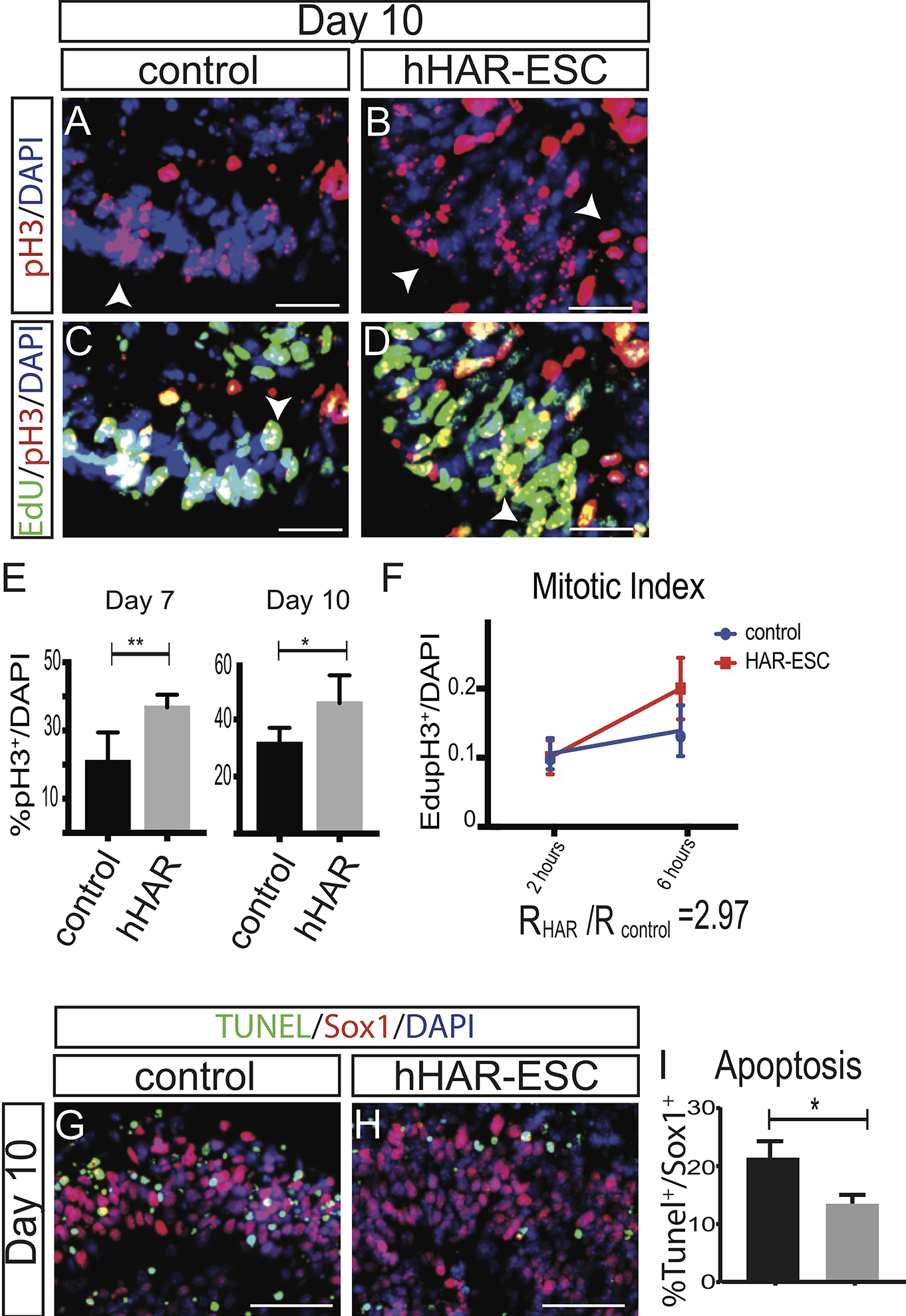
HAR-14q12 increases proliferation and decreases apoptosis in neural progenitors. A-D. Two hours EdU pulse followed by pH3 staining at day 10 in control (A,C) and hHAR-ESC (B,D) forebrain organoids indicates an increase in proliferation. Arrowheads indicate pH3^+^ progenitors. E. Quantification shows an increase in the number of pH3 positive cells in the humanized organoids at day 7 (p=0.0146) and day 10 (p=0.0453). F. Increased mitotic index in hHAR-ESC organoids calculated by comparing the amount of EdU^+^pH3^+^ cells at two time points (2 hours and 6 hours). G-I. Apoptosis (TUNEL staining) in Sox1^+^ neural progenitors is decreased in humanized hHAR-ESCs. Cell cycle and apoptosis quantifications were performed from at least 3 organoids derived from 2 independent clones. Scale bar: A-D and G-H:25 mm

In addition to cell proliferation, the final cell number can also be influenced by apoptosis, which is known to occur during normal forebrain development. However, the effect of Foxg1 on apoptosis of neural progenitors remains controversial, since both overexpression and downregulation of Foxg1 is reported to reduce apoptosis (Ahlgren *et al*, 2003; Martynoga *et al*, 2012). Upon TUNEL staining at day 10, we observed reduced levels of apoptosis in neural progenitors (Sox1^+^) derived from hHAR-ESC organoids (Fig 5G-I).

Altogether, our results suggest that the increase in Foxg1 expression, driven by the human-specific HAR-Foxg1 variant results in an expansion of forebrain neural progenitors due to increased proliferation and reduced apoptosis of neuroepithelial cells.

### Single cell RNAseq shows an expansion of radial glia and neuroepithelia progenitors in hHAR-ESC organoids

During cortical neurogenesis, human forebrain progenitors divide more often than those of other mammals, including other primates (Dehay and Kennedy, 2007). This leads not only to an increase in the final number of neurons, but also contributes to neural diversification (Wong *et al*, 2015; Dehay & Kennedy, 2007; Otani *et al*, 2016). Therefore, we wondered if the differential regulation of Foxg1 expression observed in hHAR-ESC organoids during early forebrain development impacts their cellular composition. To this end, we performed single-cell RNAseq (scRNAseq) of forebrain organoids derived from the humanized hHAR-ESCs and mouse ESC isogenic controls. Fifteen day 12 organoids of each genotype were dissociated into single cells. Similar to what was observed in the 3D organoids, hHAR-ESC samples contained 1.5x more cells than controls (data not shown). Overall, we found that the hHAR-ESCs and the mESC controls occupy contiguous, partially overlapping spaces in the t-SNE graph, indicating similar cellular identities, as expected from isogenic cell lines (Fig 6A). Clustering the cells according to their expression profiles revealed six main cell populations, which could be further divided into 23 subtypes, according with their spatio-temporal identity (Fig 6A). Five out of the six main cell types showed a neural identity, confirming the proper differentiation of the 3D forebrain organoids (Fig 6A). The remaining non-neural cluster (EnP) was a rare population (0.5% of the total cells) expressing markers of endodermal precursors, such as Sox17 and TTR.

**Fig 6.**
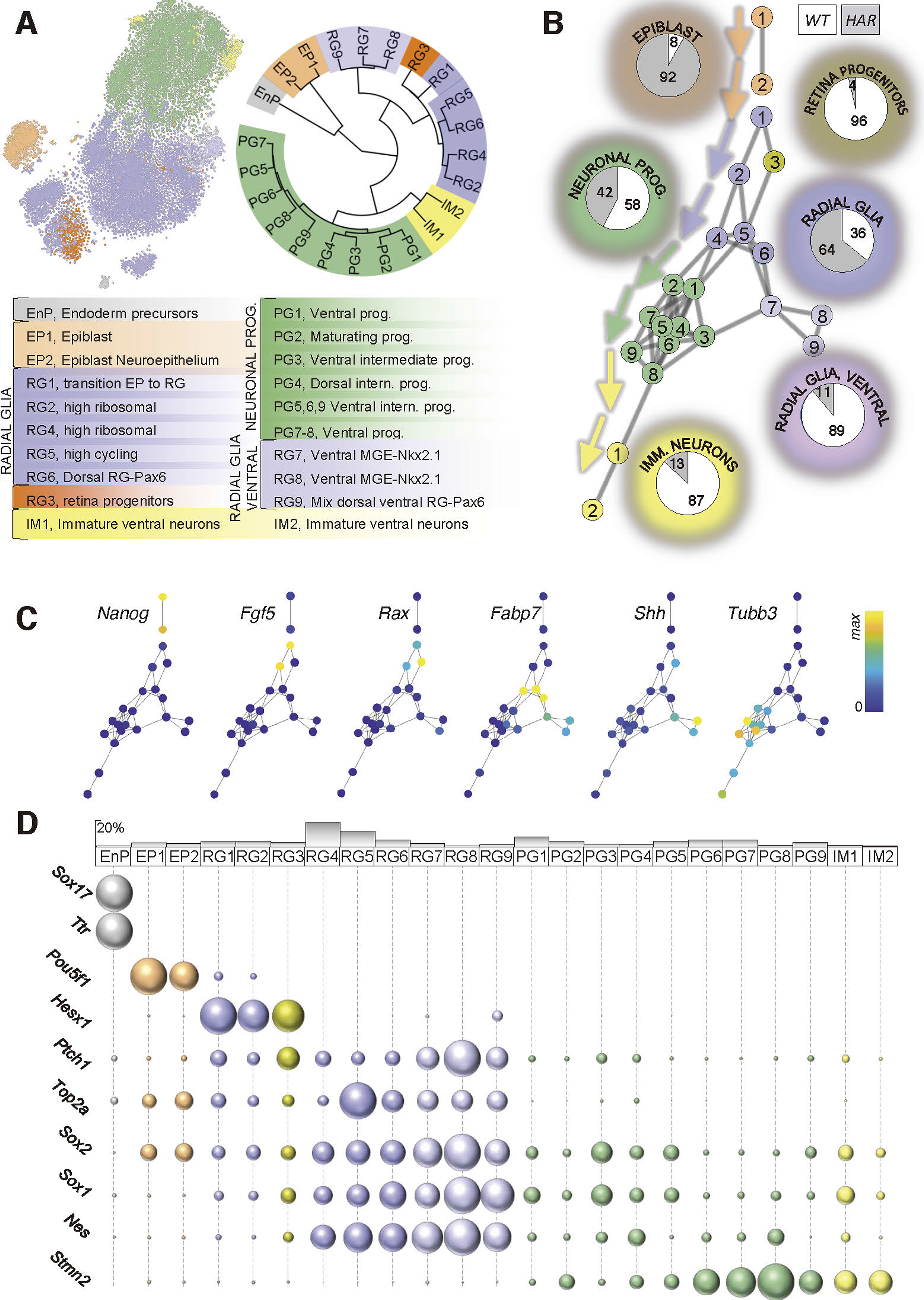
Single cell RNAseq shows an increase in neuroepithelial and radial-glia progenitors in hHAR-ESCs. A. Left: t-SNE graph showing the distribution of control (WT) and hHAR-ESC (HAR) derived organoids with a color-coded expression indicating the different subtypes of cells. Right: Dendogram representing the cluster to cluster similarity, alongside the identity of each cluster. B. Pseudotime analysis recapitulates the putative differentiation stages of the cells derived from day 12 control and hHAR-ESC organoids. Color codes of the main groups correlates with that shown in A. Pie-charts indicate the relative composition in WT and hHAR cells for each cluster group. Epiblast and RG are enriched in hHAR-ESC cells while more differentiated clusters (neuronal progenitors and immature neurons). EnP (endoderm precursors) are not represented in the pseudotime graph due to the excessive phenotypically distance from all the other cell types. The closest population to EP2 was RG1, even though not close enough to generate an edge. C. Pseudotime colored according to the average gene expression in each cluster. D. Top. Histogram representing the relative abundance of each cluster. Bottom. Detailed gene expression plots for hallmark genes.

In mouse ESC-derived 3D forebrain organoids, differentiation occurs heterochronically, meaning that progenitors at distinct developmental stages are present in the same organoid. To unravel whether there are differences in control and hHAR-ESC organoids we performed a pseudotime trajectory analysis. Each of the cell clusters was sequentially ordered according to its putative differentiation stage and correlated with its putative progeny. This analysis revealed a wide differentiation outline, ranging from undifferentiated epiblast cells/neuroepithelial (EP1,2) to immature neurons (IM1,2) (Fig 6B). Epiblast progenitors (EP2) and very primordial stages of neuroepithelial cells (EP1) were highly enriched in Sox2, Fgf5, Dpp5a and Hesx1, whilst also expressing Pou5f1 (Oct3/4) and other stem cell markers such as Nanog (Fig 6C and Fig EV4B). These cells were found in small proportions (5.8%) and are almost entirely derived from hHAR-ESC organoids (92% *vs* 8%, Fig 6B and Fig EV4). Highly proliferative radial glia cells (RG) were the most abundant cell types (55% of total cells) (Fig 6D). Our analysis identified RG with a broad range of spatio-temporal identities; starting from the most immature stages (RG1,2) still expressing low levels of neuroepithelial markers such as Hesx1, Dmnt3b and Fgf5 (Fig 6C, D, Fig EV4B), to the mature stages (RG4,5,6) expressing Fabp7. Compared to control organoids, both immature (RG1,2) and mature (RG4,5,6) telencephalic RG were overrepresented in hHAR-ESCs (overall 36% vs 64%), whereas retina RG were mainly enriched in control organoids (96% vs 4%) (Fig 6B-D, Fig EV4A). These data confirm an expansion in RG progenitors in hHAR-ESC organoids, as observed by the Nestin and Sox1 immunostainings (Fig 4N-Q). Furthermore, the hHAR-ESC organoid specific expansion of the early forebrain progenitor subpopulations (RG1,2) (Fig EV4A, S9C) was corroborated by an increase in progenitors double positive for Six3 and Otx2 in the hHAR-ESCs organoids (Fig S4A-B).

Additionally, overexpression of Foxg1 in iPSC derived from autistic patients affects the population balance between ventral (GABAergic) and dorsal (glutamatergic) forebrain neurons, leading to altered electrophysiological patterns (Mariani *et al*, 2015). Upon analyzing the patterning of the telencephalic RG, we observed that control organoids were enriched in RG populations with ventral forebrain identity (expressing Shh, Patched, Six3, Nkx2.1), whereas the humanized hHAR-ESC organoids were enriched in RGs with dorsal forebrain identity (expressing Pax6 and Emx2) (Fig 6C,D, EV4 and EV5F,H). Thus, our results hinted towards a patterning shift to dorsal telencephalic RG identities caused by hHAR activity, which was further confirmed by Nkx2.1 and Pax6 immunostaining of the organoids (Fig EV5A-D).

This differential distribution of the spatio-temporal identities of RG was also observed in more mature cell types such as neuronal progenitors (PG), and immature neurons (IM). These differentiated stages were detected in both populations; however, these more mature cell types were enriched in control organoids (Fig 6B-D and Fig EV4A-B). Moreover, the spatial patterning of PG populations showed clear commitment towards dorsal identities in hHAR-ESCs, and to ventral identities in control organoids (Fig EV5E-F).

Altogether, this analysis indicated that clusters of the most immature cells (EP and RG) were prevalent in the hHAR-ESC organoids, while control organoids showed a trend toward more advanced stages of differentiation, including neuronal/intermediate progenitors and immature neurons (Fig 6B-C). Moreover, the presence of hHAR-ESC lead to an enrichment in dorsal telencephalic fates, opposite to the ventral telencephalic signature detected more prevalently in control organoids.

These results support the hypothesis that the humanized hHAR-ESC organoids are enriched in highly proliferative early forebrain progenitors. In turn, this suggests that modifications in the early forebrain regulation by key transcription factors are likely involved in the expansion and the dorso-ventral patterning of the telencephalon in humans.

### Human enhancer variant shows functional redundancy in humans

Our analysis so far suggested that the presence of the accelerated mutations in the human enhancer variant and the resulting changes in the TFBSs may play a central role in the expansion of forebrain progenitors. To further delve into the function of these accelerated changes we explored whether mutations in the hHAR enhancer variant altered forebrain development in humans. hHAR knock-out mutations in human ESCs introduced with CRISPR/Cas9 were not viable and cells died upon transfection. Yet, we took advantage of the newly gained TFBS in the human HAR enhancer variant, namely on the acquisition of a glucocorticoid receptor (GR) binding site located in the upstream most region of the enhancer (Fig S1). We tested if we could modulate the enhancer function by using a GR inhibitor. To this end, we treated humanized mouse hHAR-ESCs and isogenic control mouse ESC-derived organoids with mifepristone (GR inhibitor) and vehicle control. The treatment was started at day 4 after induction of the neuroepithelium (Fig 7A). Indeed, GR inhibitor treatment lead to a shrinkage of the humanized hHAR-ESC organoids, as well as a decrease in Foxg1 expression, which could be still observed 5 days after stopping the treatment. These changes were not observed in control mouse ESCs (Fig 7B-G’). This indicated that the gained GR site in the human variant critically regulates Foxg1 expression and subsequent forebrain development. It also provided us with an indirect tool to understand the role of the accelerated mutations in humans. Therefore, we treated hESCs organoids with mifepristone starting at day 9; however, this treatment did not show any alteration in the organoid morphology or in Foxg1 expression (Fig 7H-J’). These different results in the activity of the human enhancer variant in mouse and humans suggest that further layers of regulation of Foxg1 expression might be operating in humans to protect this crucial developmental process.

**Fig 7.**
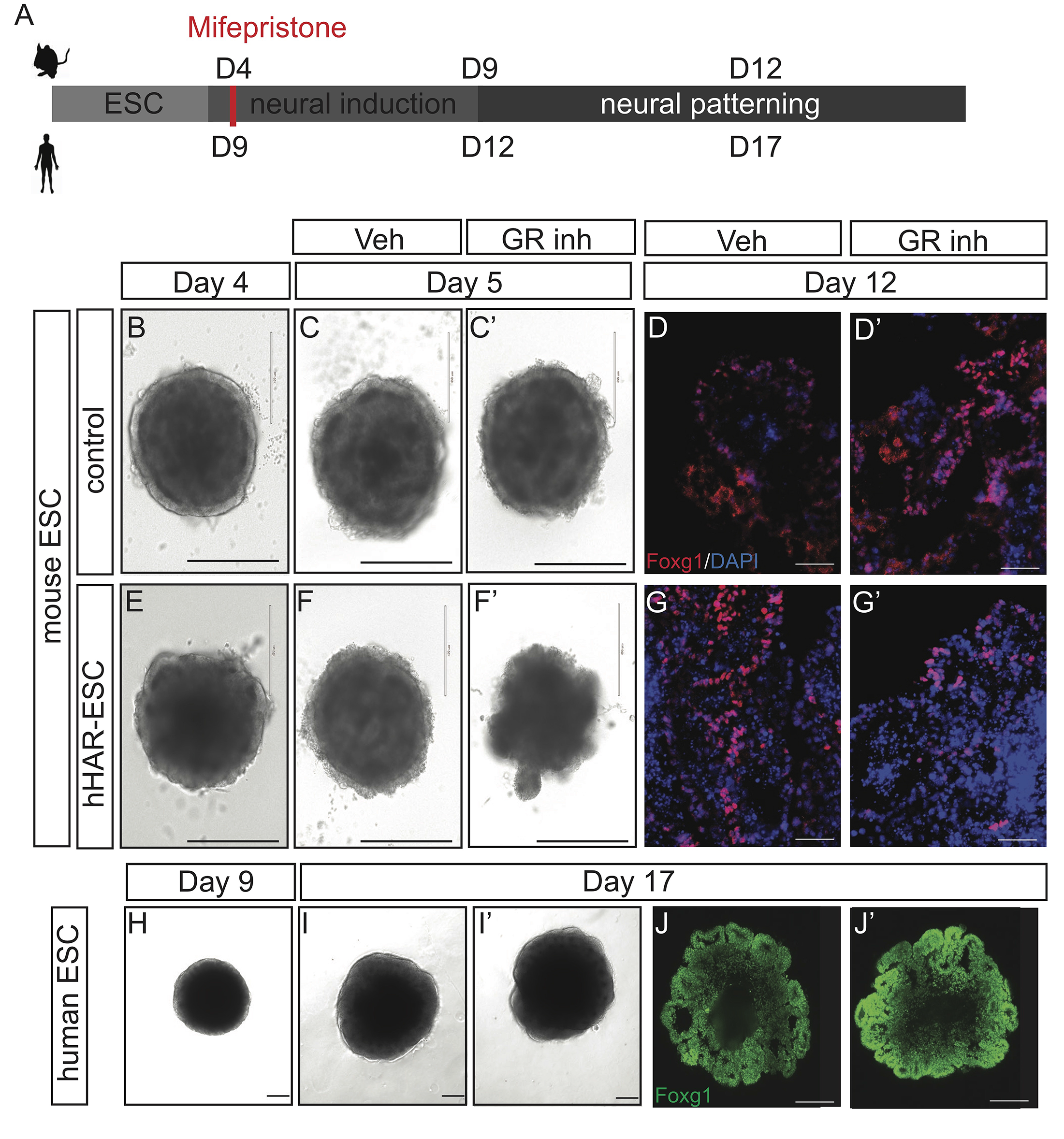
Human enhancer variant is redundant in humans. A. Scheme indicating the milestones in both mouse and human ESC-derived organoids and the time of addition of mifepristone (a GR inhibitor drug), when neural induction is starting. B-G’. mESC control (B-D’) and humanized hHAR-mouse (E-G’) under mifepristone treatment. (B,E) 24 hours after treatment, in vehicle-only (C,F) and after adding mifepristone (C’-F’) showing a disruption in the differentiation of the humanized organoids. D,D’ and G,G’. Reduction in Foxg1 expression after treating with mifepristone is only detected in humanized organoids at day 7. H-J’. Human ESC brain organoids showing no effect upon the administration of mifepristone from day 9-17 neither in morphology nor in the expression of Foxg1. This suggests that the presence of mutations in heterozygosity in this enhancer might not be enough to be pathological, suggesting certain level of redundancy in the enhancer function. Scale bars: B,C,C’,E,F,F’:400mm; D,D’,G,G’, H-J’: 100mm

## Discussion

Minor changes during early embryonic stages can lead to critical developmental alterations. Most likely, this is one of the main reasons why molecular and morphological features are highly conserved among different organisms during early embryonic stages (gastrula to neurula) (Sasai & De Robertis, 1997). However, despite this conservation, few but important changes are responsible for the evolutionary divergence among species. This is particularly evident in the case of the brain, where the expansion of the human forebrain is one of the most prominent and rapid evolutionary changes among mammals. The evolutionary genetic changes involved in the expansion of the forebrain encompass two main mechanisms: the generation of novel genes through gene duplication (Fiddes *et al*, 2018; Suzuki *et al*, 2018; Charrier *et al*, 2012; Florio *et al*, 2015), and differential gene regulation at the transcriptional (i.e., enhancers) (Boyd *et al*, 2015) and post-transcriptional (i.e., lnc-RNAs) (Pollard *et al*, 2006) levels. In this study, we present new evidences about the functional role of human-specific substitutions in an active ultraconserved enhancer controlling Foxg1 expression during early neural development stages and their consequences on human forebrain evolution. We found that the human variant of this enhancer (HAR-14q12) shows increased enhancer activity when compared with its mouse or chimp counterparts, resulting in increased levels of Foxg1 expression. In addition, gain-of-function experiments, either in transgenic HAR-14q12 mice or targeted humanized mouse ESC-derived 3D forebrain organoids resulted in the expansion of neuroepithelial and radial glia progenitors. Our results provide an additional mechanism that has potentially contributed to the shorter cell cycle and higher proliferative potential of RG progenitors, which is proposed to drive the evolutionary expansion of the mammalian telencephalon (Calegari *et al*, 2005; Pilz *et al*, 2013; Betizeau *et al*, 2013; Pilaz *et al*, 2009; Reillo *et al*, 2017).

Foxg1 is one of the earliest genes to be expressed during neural patterning and its expression levels are highly dynamic during forebrain development. High levels of Foxg1 in the ventral telencephalon are key for the correct formation of the ventral midline; however, later during development, the dorsal telencephalic expression is higher than the ventral (Geng *et al*, 2016). Our data shows that the humanized forebrain organoids have an expansion in the number of dorsal telencephalon progenitors. This could be a consequence of the direct regulation of Pax6 by Foxg1 (Manuel *et al*, 2011) during early stages, favoring eventually the expansion of cortical progenitors.

Enhancer redundancy during development is a protection mechanism for the organisms to ensure the robustness of critical processes (Osterwalder *et al*, 2018; Pennacchio *et al*, 2006). We approached this question investigating the presence of pathogenic mutations in epileptic and FOXG1 syndrome patients. We detected a slight increase in the presence of mutations in heterozygosity in patients than in the control cohort (data not shown). Similarly, the different phenotypes observed when the HAR enhancer was chemically knocked-down using humanized mouse- and human-ESCs suggest that the human enhancer might be redundant in its function during critical steps of Foxg1 expression in brain development.

Altogether, this study shows for the first time the critical role that human-specific nucleotide substitutions in regulatory regions of key developmental genes play during early embryonic stages in the expansion and patterning of the telencephalon through the specific enlargement of neural progenitors and RG cells.

## Methods

### Isolation of early stage mouse embryos

Inbred NMRI females were mated and litters harvested at E7.5-8.0. Embryos were kept on ice and processed directly or frozen at −80C in 20 ml of RNAse/DNAse free PBS supplemented with 1× protein inhibitor (Diagenode, Belgium). Embryos younger than late gastrula (younger than E7.5) and older (more than 4-6 somites) were discarded. These procedures were performed according the local IACUC regulations.

### H3K27ac ChIP-seq

Three E7.5-E8 litters were pooled together and embryos were dissociated mechanically using a P20 tip. ChIP was performed using the Low Cell ChIP kit (Diagenode, Belgium). Briefly, DNA was crosslinked in formaldehyde and sonicated in a Bioruptor for 15 minutes at high power (Diagenode). Cells were resuspended in a total volume of 250 ml of Buffer A and Buffer B mix (half of the recommended volume for 0.5×10^6^cells). ChIP was performed using Protein G beads (Diagenode) O/N using a H3K27ac antibody (Active Motif, Ref:39135). Immunoprecipitated DNA was diluted using iPURE kit (Diagenode). DNA yield and quantity were analyzed in a Bioanalyzer immediately and kept at −80c until NGS was performed. Three replicates containing 3 litters were analyzed. From E8.5 embryos, anterior head and tails were isolated and from E12.5 embryos forebrain and tails were dissected. Tails were used as negative controls. DNA was resuspended using Low Cell Kit (Diagenode) and RT-PCR was performed using Sybr Green Mix (Applied Biosystems). Primer used for qPCR forward 5’- GGCCCTTAGCACTCTCTTTG-3’ and reverse 5’- TGGGAGCTGGTATTTTTGGA-3’. The database is publicly available in GEO with number GSE116071.

### ChIP-Seq data processing

Initially BWA (version 0.5.9-r16, default parameter) was used to align the ChIP-Seq reads to the mouse genome mm9 (MGSCv37 from Sanger) after trimming adapters by cutadapt (version 1.13), biobambam (version 2.0.57), then they have been used for marking duplicated reads. Only non-duplicated reads have been kept by SAMtools software (parameter “-q 1 -F 1024” version 1.4). We followed the ENCODE criteria for quality control (QC) on the data. Non-duplicated version of SPP (version 1.11) has been used to draw cross-correlation and calculated relative strand correlation value (RSC) under support of R (version 2.14.0) with packages caTools (version 1.17) and bitops (version 1.0-6) and estimated the fragment size. After confirmation, replicates have RSC > 1, data was merged and peaks were called with MACEV1 (version 2.0.9 20111102, option “nomodel” with “extsize” defined as fragment size estimated by merged data as above) with FDR corrected p-value cutoff 0.05 (strong peaks) or 0.5 (weak peaks). Extended reads at the called peaks were counted and the consistency between replicates confirmed. Extended reads were pooled and used to generate the final bigwig track for visualization.

To check whether they overlap with HARs, we liftOver reads from mm9 to hg19 using CrossMap (version 0.1.5) and generated bedGraph files normalized to 15M total reads by bedtools (version 2.24.0). Then converted to bigwig format by bedGraphToBigWig (UCSC version 4). For the other tracks (Fig 1F), we downloaded the bigwig files from ENCODE and liftOver them to hg19 also by CrossMap if they are mm9.

Selection of genes was performed using the annotation in gene ontology database as brain development (GO:0007420), nervous system development (GO:0007399), development (GO:0032502) (gene lists extracted by R package either from human org.Hs.eg.db or mouse org.Mm.eg.db). Only 5 HARs have been found within 1kb of strong peaks, and 766 genes within 1Mb of the HARs.

### Molecular Phylogenetic analysis by Maximum Likelihood method

Sequences of orthologous HAR_755 region from selected species have been extracted from MAF file download from UCSC as multiple alignments of 100 vertebrate species(hg19) and complete aligned by ClustalX 2.1 using default parameters. The evolutionary history was inferred by using the Maximum Likelihood method based on the Tamura-Nei model (Tamura and Nei, 1993). The tree with the highest log likelihood (−1720.38) is shown. Initial tree(s) for the heuristic search were obtained automatically by applying Neighbor-Join and BioNJ algorithms to a matrix of pairwise distances estimated using the Maximum Composite Likelihood (MCL) approach, and then selecting the topology with superior log likelihood value. The tree is drawn to scale, with branch lengths measured in the number of substitutions per site (Fig 1D, numbers next to the branches). The analysis involved 10 nucleotide sequences. All positions containing gaps and missing data were eliminated. There was a total of 777 positions in the final dataset. Evolutionary analyses were conducted in MEGA7(Kumar *et al*, 2016).

### LacZ reporter transgenic mice

The human variant of the enhancer was amplified from human DNA (derived from the 293T cell line) using specific primers, designed using the SnapGene software, and inserted into a hsp68:LacZvector (Pennacchio *et al*, 2006) by Gibson Assembly. The generated HAR-Foxg1-LacZ vector was injected by pronuclear injections and transferred into stimulated females. Embryos were dissected 9 days and 12.5 days after transfer. E9.5 and E12.5 were fixed in glutaraldehyde and paraformaldehyde mix freshly prepared for 20 mins to 3 hours. X-gal staining was then performed and monitored under the scope. E9.5 and E12.5 embryos were stained for 2-3 hours to one day at 30°C. X-gal reaction was stopped by fixing the embryos in paraformaldehyde and cleared in glycerol prior to imaging in a Leica Steroscope Microscope. After dissection, some embryos were fixed in paraformaldehyde and embedded in OCT. They were kept at −80C in methanol until b-galactosidase immunostaining coupled with Foxg1 ISH or eosin staining. For further details see section below.

### hHAR-Foxg1 and mHAR-Foxg1 transgenic mice

Foxg1-GFP vector (Baek *et al*, 2015) was obtained at Addgene. Mouse or human enhancer variant were amplified similarly and introduced in the donor vector coding for Foxg1-GFP. Transgenic embryos with hHAR::Foxg1-GFP or mHAR::Foxg1-GFP were generated by pronuclear injection and transferred into stimulated females. Embryos were harvested at E9.5, fixed in paraformaldehyde 4% 2 hours, cryopreserved in sucrose and embedded in OCT. 10 mm serial sagittal cryosections were stained for immunofluorescence.

### Luciferase analysis

Human (hHAR), mouse (mHAR) and chimp (ptHAR) enhancer variants were amplified from 293T cell line, W9.5 ESCs, and chimp (kindly provided by Dr. Debra Silver, Duke University, NC, USA) DNA. The hHARshort fragment (Fig EV1G) was generated by HindIII restriction of the full-length PCR fragment. PCR fragments were cloned into the pGL4.10 vector. 100 ng of each vector was transfected into 293T cells using Lipofectamine 2000 (Promega). Empty renilla vector was used as negative control (1ng/well). Briefly, 293T cells were passaged twice after thawing and the night before were plated at 60-70% confluency in 24-well plates. Next morning, transfections were performed and cells were kept in Opti-MEM for 24 hours. After that, Luciferase activity was measured with the Dual-Glo kit (Promega). pGL4.10 empty vector was used as normalization value. Three experiments in quadruplicates were performed. ANOVA was used for statistical analysis.

### Generation of mHAR^ko^/hHAR Foxg1:Venus ESC

The Foxg1-Venus ESCs were kindly provided by Dr. Sasai (CDB, Riken, Japan). CRISPR/Cas9 was used to generate a heterozygous homologous recombination (HR) with hHAR-Foxg1 and indel mutations in the second allele as described elsewhere (Acosta *et al*, 2018). Briefly, the human enhancer variant flanked by 700 bp homology arms (5’ and 3’ to the enhancer) to the mouse endogenous locus were cloned into a pBSK backbone vector. Donor vector and gRNA-containing vector specific for the locus were transfected into Foxg1-Venus ESCs. Forty eight hours later, cells were FACS sorted for Orange Fluorescent protein and seeded on feeder layers. 48 colonies were picked and analyzed for the presence of indel mutations using T7E1 endonuclease analysis. The presence of the human variant in heterozygosis was confirmed through Sanger sequencing. ESCs were maintained as reported in maintenance media with blasticidin and LIF (Eiraku *et al*, 2008).

### Humanized hHAR-Foxg1 ESCs

W9.5 wild-type mouse ESCs were targeted in homozygosis using the vector carrying the human enhancer variant flanked by 700 bp homology arms at the 5’ and 3’ ends. Two gRNA were located in the flanking region of the mouse enhancer variant were designed using the http://crispr.mit.edu website. The gRNAs were cloned into GeneArt OFP vectors (Life Technologies) and bi-allelic homologous recombination was performed as described (Acosta *et al*, 2018). Briefly, cells were transfected with the 1mg of the donor vector and 500 ng of each of the gRNA vectors using Amaxa Nucleofector (Lonza) and the predetermined program for mESC (A23). Cells were treated for 48 hours with the NHEJ repair inhibitor SCR7 and clones were picked and expanded. Fourteen clones were identified as positive for the presence of the human enhancer variant in homozygosis. Three independent clones were chosen to perform the experiments described (#4, #7 and #9). ESC lines were maintained in ES Media on feeder layers. Cells become Feeder-free culture upon three passages on 2i media. The presence of off-target was analyzed for the most common locations for each of the gRNAs using T7E1 analysis as reported in our previous article (Acosta *et al*, 2018).

### Mouse ESC-derived 3D organoids

ESCs were thawed and maintained in ES media (W9.5 isogenic ESCs) or MM media (Foxg1-Venus ESC) for at least three passages before starting the differentiation. 3D ESC-derived organoids were differentiated following a modified version of Sasai’s group protocol (Nasu *et al*, 2012). Briefly, feeder-free ESCs were trypsinized and approximately 3500 cells were plated onto low-adherence plates (Nunc) in chemically-defined media (CDM) and 2-2.5% Matrigel as extracellular matrix (Corning).CDM was prepared as follows: Iscove’s modified Dulbecco’s medium (IMDM; invitrogen)/Ham’s F12 medium (invitrogen) 1∶1, 1× Chemically-defined lipid concentrate (invitrogen), 450 μM monothioglycerol (Sigma), 5 mg/ml purified bovine serum albumin (>99% purified by crystallization; Sigma) and 15 μg/ml apo-transferrin (Sigma). At day 7 of differentiation, 3D organoids were transferred to a bacteria-culture plate and maturation media in hyperoxia conditions (40% O_2_). Media was replaced every other day until day 12 (Fig 3A). If the organoids were to stay up to day 25 in culture, media was substituted by N2:B27 media (Gaspard *et al*, 2008). Organoids were collected with a wide-gauge pipette tip and transferred in the required solution for subsequent analysis.

Treatment with mifepristone (GR inhibitor) was performed from day 4 to day 7 at a concentration of 200 nM.

### Human embryonic stem cell culture

Feeder free H9 Human ESCs were obtained from WiCell and verified for normal karyotype. They were cultured in a feeder free manner on Matrigel (Corning; hESC-qualified matrix) coated plates using mTeSR medium (Stemcell technologies) in a 5% CO_2_ incubator at 37°C. Cells were routinely checked for genomic integrity and contamination and tested negative for mycoplasma.

### Human ESC-derived organoids

Human cerebral organoids were cultured using minor modifications of the previous protocol (Lancaster *et al*, 2016). Briefly, to generate embryoid bodies (EBs), hESCs colonies were dissociated into single cell suspension using accutase (Stem cell technologies). 9000 cells per well were seeded in an ultra-low attachment round bottom 96-well plate (Corning) in mTesr medium (Stem cell technologies), supplied with 50μM Rho-associated kinase (ROCK) inhibitor (Calbiochem). EBs were fed on day 3 and 5 with mTesr medium. From day 6 onwards EBs were cultured in neural induction medium to form neuroepithelial tissue with daily replenishment of the medium. On day 12, EBs were embedded in Matrigel (Corning) droplets and cultured in a differentiation medium without vitamin A and the medium was replenished every two days.

The drug treatment was performed from day 9 onwards and continued till day 17. Mifepristone stocks were diluted to 200μM aliquots and used at 200nM concentration with equivalent amount of ethanol as a vehicle control.

For immunostainings, on day 17, the organoids were treated with cell recovery solution (Corning) to remove the Matrigel and then fixed with PFA. The fixed organoids were later cryosectioned and stained to visualize Foxg1 expression using (Abcam ab18259 antibody).

### Immunostaining and in situ hybridizations

Samples, either derived from ESC-organoids or from mice, were fixed in paraformaldehyde at 4°C from 15 minutes to O/N depending on size. Next, they were embedded in OCT and sectioned 10-14 μm thick and kept frozen (−20C) until used. For immunostaining, cells were thawed and blocked with 5% BSA, 3% Horse or Bovine serum, 0.1% Triton (unless specified-differently) in PBS or TBS. Primary antibodies were incubated O/N at 4°C in a wet chamber. After washing in PBS-0.1%Triton, sections were incubated with secondary antibodies for 1 hour at RT and mounted with Fluoromount. Images were acquired with a Zeiss AxioCam epifluorescent microscope or a Leica Confocal Microscope. In situ hybridization were performed as described previously (Geng *et al*, 2016).

### Gene expression qPCR

RNA was extracted with the RNAeasy kit (Qiagen) and retrotranscription was performed with M-MLV retrotranscriptase (Clontech). Subsequently, qPCR was performed using Sybr Green Master Mix (Applied Biosystems). Primers used were: Emx1 F:5’GAGCGAGCCTTTGAGAAGAA3’; R:5’CCAGCTTCTGCCGTTTGTAT3’; Foxg1 F:5’TGGGAGATAGGAAAGAGGTG3’; R:5’GTGGTGGTGATGATGATGG3’

### Mitosis measurements

For the mitotic index evaluation, a pulse of EdU was performed in 3D organoids using the Click-it EdU kit (ThermoFisher). Organoids were collected 2 and 6 hours after the EdU pulse for the pulse-and-chase experiment. Cells were quantified using Image J, counting the total DAPI, EdU and pH3 cells for each layer.

### Cell Cycle Flow cytometry

Cell cycle was determined in ESCs to verify they behave similarly to controls. Briefly, isogenic controls and hHAR-ESCs were thawed in feeder-free conditions and maintained in culture for 3 passages. Subsequently, they were trypsinized and Propidium Iodide was added to the cell solution and incubated at 4°C for 30 minutes. Cell cycle was determined using Fortessa Flow Cytometre (BD).

### Single-cell RNAseq analysis

Single cell suspension was prepared from 15 day 12 control or humanized hHAR-ESC organoids using a TryPLE (Gibson, US) solution complemented with DNAse and ROCK inhibitor. Viability exceeded 90% as determined by AO/PI staining on the Nexcelom K2 Cellometer. Single cell RNA-seq libraries were prepared using Chromium single cell controller and 3’ V2 reagents (10× Genomics) according to standard protocols. Libraries were sequenced on an Illumina HiSeq 4000 instrument, one library per lane. Data were processed using Cell Ranger V2.0 pipeline with standard parameters (10× Genomics) and reads were mapped to mm10 reference genome. Overall, 14123 cells (5636 in control *vs*8414 cells in hHAR-ESC) satisfied these quality requirements and were used for the subsequent analysis. To cluster single cells, detect markers and estimate pseudotime differentiation the tool bigSCale was used (Iacono *et al*, 2018). bigSCale is an analytical framework for large scale single-cell data analysis and generates a heuristic model of the noise and sparsity of scRNA-seq data, ultimately allowing to cluster individual cells according to their real phenotypic similarity. The use of iCells, a bigSCale feature allowing the analysis of datasets >50k cells, was not needed for our dataset. Initially, a clustering of the whole dataset revealed a number of well separated clusters plus two large groups that we recognized as neuronal progenitors and radial glia. To achieve a better characterization of the subtypes of the latter populations, we next re-clustered them individually, reaching a final number of 23 final clusters. The latest version of bigSCale also allows to generate a pseudotime trajectory which is based on a graph approximation of cluster to cluster phenotypic distances. The databases will be publicly available and the GEO numbers will be provided at the time of publication.

## Acknowledgements

We would like to thank Geoffrey Neale from St Jude Children’s Research Hospital for the NGS sequencing advice on H3K27ac ChIP-seq, Rajesh Awatramani and Lynn Doglio at the NU Transgenic Core Facility and Linda Degenstein at University of Chicago for the generation of transgenic and chimeric mice and Paul Mehl at the NU Flow Cytometry Core Facility for FACS sorting. We thank Nelle Lambert for valuable comments on the manuscript. This work was partially supported by a grant of the National Institutes of Health (EY12162) to G.O. S.A. was funded by the International FOXG1 Foundation and Beatriu de Pinós Fellowship program (grant 2017-BP-00176) by Generalitat de Catalunya. H.H. is a Miguel Servet CP14/00229) researcher funded by the Spanish Institute of Health Carlos III (ISCIII). A.V. M is funded by NIH grants HL135124 and AI135964 and Department of Defense grant PR141319.IR and MM were funded by the Spanish government (grant BFU2017-84914-P), and work at the CNIC is supported by the Institute of Health Carlos III (ISCIII), the Ministerio de Ciencia, Innovación y Universidades (MCNU), the Pro CNIC Foundation, and is a Severo Ochoa Center of Excellence (SEV-2015-0505).

## Author contributions

S.A. and G.O. designed the study and wrote the manuscript with input from the remaining authors. S.A., L.F., N.T. performed most of the experiments, including the 3D organoids model and the transgenic mice analysis. J.S and J.K. designed and performed the hESC experiments. I.R. and M.M. performed the LacZ transgenic embryos analysis. G.I., A.V.M., N.J. and H.H. performed and analyzed single cell RNA-seq. B.X. performed the bioinformatics analysis on ChIP-seq and HAR databases. M.S. and E.W. analyzed the ancient hominin databases.

